# MitoToxy Assay: a novel cell-based method for the assessment of metabolic toxicity in a multiwell plate format using a lactate FRET nanosensor, Laconic

**DOI:** 10.1101/583096

**Authors:** Y Contreras-Baeza, S Ceballo, R Arce-Molina, PY Sandoval, K Alegría, L.F. Barros, A. San Martín

## Abstract

Mitochondrial toxicity is a primary source of pre-clinical drug attrition. Methods that detect mitochondrial toxicity as early as possible during the drug development process are required. Here we introduce a new method for detecting mitochondrial toxicity based on MDA-MB-231 cells stably expressing the genetically-encoded FRET lactate indicator, Laconic. The method takes advantage of the high cytosolic lactate accumulation observed during mitochondrial stress, regardless of the specific toxicity mechanism, explained by compensatory glycolytic activation. IC_50_ determination using a standard multi-well plate reader, shown that the methodology allowed to detect metabolic toxicity induced by azide, antimycin, oligomycin, rotenone and myxothiazol with high sensitivity. Suitability for high-throughput screening applications was evaluated resulting in a Z’-factor > 0.5 and CV% < 20. A pilot screening allowed sensitive detection of commercial drugs that were previously withdrawn from the market due to liver/cardiac toxicity issues, such as camptothecin, ciglitazone, troglitazone, rosiglitazone, and terfenadine, in ten minutes. We envisage that the availability of this technology, based on a fluorescent genetically-encoded indicator will allow direct assessment of mitochondrial metabolism, and will make the early detection of mitochondrial toxicity in the drug development process possible, saving time and resources.

## Introduction

High-throughput screening (HTS) is a fundamental step in the hierarchical and long drug development process. Discovering and developing a drug starts with libraries consisting of thousands of chemicals, which are screened for a target activity using *in vitro* assays. Hits are selected for subsequent stages, then tested for activity and toxicity in cells, tissues, animal models, followed by clinical trials. Most candidates fail in clinical trials, as they are found to be ineffective in a physiological context or produce unwanted side effects. Rate of failure of lead compounds during the drug development process is called attrition, which can be quantified giving a wide view about the whole efficacy of drug development chain.

Pre-clinical and clinical safety, pharmacokinetics or bioavailability issues, and rationalization of the company portfolio are the main sources of drug attrition (Waring et al. 2015). However, toxicity-related side-effects are by far the most important source of attrition during pre-clinical stages, being responsible for 59% of failures (Waring et al. 2015). For instance, a comprehensive study combined data on drug attrition from AstraZeneca, Eli Lilly, GlaxoSmithKline, and Pfizer, which revealed that during 2000 and 2010 a total of 211 from 356 lead compounds were rejected due to safety concerns in the pre-clinical stage (Waring et al. 2015). Attrition at later stages of the development chain produces a higher economic impact and is the most important determinant of overall efficiency. These data suggest that although minimizing safety-related attrition has been a significant area of investment across the industry, it remains a key area to be improved.

Mitochondria is the main source of toxicity issues during drug discovery campaigns and clinical trials (Dykens and Will 2007; Nadanaciva and Will 2011; Vuda and Kamath 2016; Wallace 2008; Wallace 2015). For instance, more than 80 individual chemical identities that inhibit complex I of the Electron transport chain (ETC) alone have been reported (Dykens and Will 2007). Potential mitochondrial targets of drug-induced toxicity are not only restricted to primary targets such as ETC or uncoupled mitochondrial oxidative phosphorylation, but also to secondary targets including molecular regulation, substrate availability, and protein trafficking (Wallace 2015). Higher numbers are expected due to the highly diverse mitochondrial proteome and the complexity of mitochondrial physiology.

Current methodologies to evaluate mitochondria damage take advantage of: i) *in vitro, in cellulo, ex vivo* and *in vivo* systems; ii) samples such as isolated mitochondria, permeabilized cells, living cells, tissue and animal samples and iii) readouts such as mitochondrial membrane potential (MMP), oxygen consumption rate (OCR), Adenosine Triphosphate (ATP), extracellular acidification rate (ECAR), Reactive Oxygen Species (ROS), Glutathione (GSH), and viability assays. Also, a computational method to predict *in vivo* toxicity was recently described (Huang et al. 2016). The ideal method for pharmaceutical industry has to offer a balance between throughput and physiological relevance. For instance, *in vivo* models provide low throughput because the sample preparation for measurements are laborious, but the physiological relevance is high because the whole animal presents intact short and long-distance interactions between molecules, cells, and organs that allow the testing of key parameter such as bioavailability and cellular toxicity of lead molecules. Additionally, *in vitro* methodologies such as isolated mitochondria offer high-throughput with low physiological relevance since they are easy to set up in a multi-plate reader, but lack cytosolic, cell-to-cell, and organ context. New technologies are needed to complement the current methodologies and achieve a balance between throughput and physiological relevance.

Genetically-encoded fluorescent indicators provide a minimally invasive approach for metabolite detection in living systems (Okumoto et al. 2012; San Martin et al. 2014b). They are built by the fusion of a ligand-binding moiety from bacteria to a fluorescent protein FRET (<u>F</u>oster <u>R</u>esonance <u>E</u>nergy <u>T</u>ransfer) pair module. Binding of the test molecule, triggers a conformational change that affects the relative distance and/or orientation between the acceptor/donor fluorescent proteins, causing an increase or decrease in FRET efficiency (Miyawaki et al. 1997). These genetically-encoded indicators have been instrumental to decipher molecular mechanism involved in fast modulation of oxidative and glycolytic metabolism in the brain (Baeza-Lehnert et al. 2018; Fernandez-Moncada et al. 2018; Lerchundi et al. 2015; Machler et al. 2016; San Martin et al. 2017; San Martin et al. 2014b; Valdebenito et al. 2016). Being fluorescent and genetically-encoded, these tools have a great potential to develop cell-based high-throughput methods for the pharmaceutical industry (Mendelsohn et al. 2018; Schaaf et al. 2018; Schaaf et al. 2017; Zhao et al. 2015; Zhao et al. 2018). Using a FRET-based lactate indicator Laconic (San Martin et al. 2013), we have observed that the mitochondrial damage induced by sodium azide or physiological mitochondrial modulator signals such as nitric oxide, produces a fast and acute cytosolic lactate accumulation (San Martin et al. 2017; San Martin et al. 2013). A phenomenon explained primarily by a reduction of mitochondrial pyruvate consumption and secondary glycolysis activation (Bittner et al. 2010; San Martin et al. 2017; San Martin et al. 2014a). Both effects converge into increased cytosolic pyruvate and NADH levels, displacing the chemical equilibrium of LDH (Lactate Dehydrogenase) to lactate production within seconds. Therefore, we hypothesized that cytosolic lactate is a fast and sensitive metabolic beacon of mitochondrial dysfunction.

This paper describes a cell-based method to detect metabolic toxicity by measuring cytosolic lactate accumulation induced by mitochondrial damage. To develop and implement the methodology, we generated a cancer cell line MDA-MB-231 that stably expresses Laconic. To assess the suitability for high-throughput screening applications, Z’-Factor and inter-plate variability were determined using classical mitochondrial toxicants. As a proof of the concept, 13 compounds were tested in a pilot screening. The method provided rapid detection in 10 minutes of a panel of classical mitochondrial toxicants and allow the detection of previously described toxic drugs in a 96-well format using standard multi-plate readers.

## Materials and Methods

Standard reagents and inhibitors were acquired from Sigma, Tocris and Toronto Chemicals. Plasmid encoding FRET sensor Laconic is available from addgene (www.addgene.org). Ad-Laconic (serotype 5) was custom made by Vector Biolabs. LV_Laconic was generated in CECs using Viral Power System from ThermoFisher. MDA-MB-231 cell line was acquired from ATCC (ATCC^®^ CRM-HTB-26™).

Mitochondrial toxicant stocks were prepared as follow: Azide was dissolved in distilled water, antimycin in 0.16% ethanol, myxothiazol, oligomycin, and rotenone in 0.8% DMSO (Dimethyl Sulfoxide). The 13 compounds used in the pilot screening were dissolved in DMSO. Maximal final concentration of 0.2% DMSO. Master stock of pCMBs at 250 mM (5000x) was prepared in 10 M NaOH.

## Brain Cell Culture

All animal procedures for the *in vitro* experiments were approved by the Institutional Animal Care and Use Committee of the Centro de Estudios Científicos. Animals used for primary cultures were mixed F1 male mice (C57BL/6J × CBA/J), kept in an animal room under SPF conditions at a room temperature of 20 ± 2 °C, in a 12/12-h light/dark cycle and with free access to water and food. Mixed cortical cultures of neuronal and glia cells (1 – to 3-day-old neonates) was cultured in Neurobasal medium (Gibco) supplemented with 5 mM glucose and B27. Cells were seeded on petri dishes and superfused with KRH HEPES pH 7.2 equilibrated with 21% air, 5% CO^2^. The lactate imaging experiments with single-cell resolution were performed using cultured astrocytes at 11 – 14 days post-dissection. Real time experiments were performed with KRH HEPES bathing solution containing (in mM) 136 NaCl, 3 KCl, 1.25 CaCl_2_, 1.25 MgCl_2_, 10 HEPES/Tris and pH 7.2 at 36°C with 300 mOsm, without carbon sources for mitochondrial toxicant experiments and 2 mM glucose and 1 mM lactate for Warburg index determination. Experiments were performed in an upright Olympus FV1000 confocal microscope equipped with a 20X water-immersion objective (numerical aperture, 1.0) and a 440 nm solid-state laser.

## Generation of a Clonal Laconic Cell Line

Recombinant lentiviral particles for Laconic were produced by co-transfection of lentiviral vectors LV-Laconic and ViralPower^®^ lentiviral packaging mix (ThermoFisher) in HEK293FT cells. Viral supernatants were harvested 72 hours after transfection, filtered through 0.45 μm units (Millipore), concentrated and store at −80°C. Transductions with Laconic lentivirus (1.3×10^5^ TU/ml) were performed at a MOI (<u>M</u>ultiplicity <u>o</u>f <u>I</u>nfection) of 3 in MDA-MB-231 cells. The selection of stable sensor expression was established under selective pressure using Blasticidin at 5ug/ml for 2 weeks after transfection. Cell cultures were processed by FACS (<u>F</u>luorescence-<u>A</u>ctivated <u>C</u>ell <u>S</u>orting) in order to isolate clonal cell lines of MDA-MB-231 expressing Laconic. Highly fluorescent cells were sorted, and single cells were collected individually in a 96 well plate. Each clone was propagated and screened based on the expression of the functional Laconic sensor.

## MitoTox Reporter Cell Culture and Measurements

MitoTox reporter cell line was cultured in Leibowitz media (Gibco) supplemented with 10% fetal bovine serum at 37 °C without CO_2_ equilibrated atmosphere in a 100 mm petri dish. Once cells reached 100% confluency, they were trypsinized using 0.5% trypsin-EDTA (Gibco) and washed in PBS. Viability was determined using trypan blue staining and 10,000 cells were seeded into each well of a 96 well plate. Experiments in 96 well format were performed once cells reached 100% confluency after 6 days, during incubation the media was not replaced. All the plates with signal-to-noise ratio below 1.3 for mTFP and Venus channels were discarded. Background signal was determined with KRH buffer without cells.

Experiments consisted in washing out the culture media using KRH HEPES buffer containing (mM) 136 NaCl, 3 KCl, 1.25 CaCl_2_ 1.25 MgCl_2_, 10 HEPES/Tris and pH 7.2 at 37°C with 300 mOsm. Then 200 μl of KRH HEPES buffer without any carbon sources was added to each well and the baseline was measured (R_MIN_), next 100 μl of buffer was replaced with mitochondrial toxicant/controls and signal was measured to obtain a stimulated response (R_MAX_). In experiments where pCMBs was used to block lactate exit, the inhibitor was added simultaneously with the mitochondrial toxicants. Data was collected in triplicate after 5, 10, 30, and 60 minutes of incubation. All the experiments in 96 well plates were performed in an EnVision^®^ multiplate reader (PerkinElmer). Each well was excited at 430 nm and the intensity of mTFP and Venus fluorescence emissions were recorded at 485 nm and 528 nm, using a set of filters.

## Data Analysis

To plot the data, we used a ratio between mTFP and Venus channels. This ratio is proportional to lactate levels, since its binding to the sensor produced a decrease in FRET efficiency. This ratio was expressed in two formulas described below:

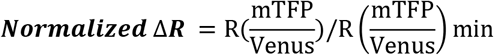

Where R(mTFP/Venus)min corresponded to the minimal change of mTFP/Venus ratio during the experiment. Results from experiments in 96 well plates were expressed by the ΔR%. This parameter was obtained from the difference between the R_MAX_ and R_MIN_ from the same well (before-after experiment) as follow:

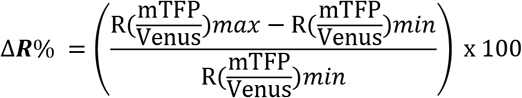

To validate the suitability of the assay for high-throughput screening a coefficient called Z’-factor was calculated (Zhang et al. 1999). This coefficient is reflective of the assay signal dynamic range and the data variation associated with the signal measurements. The Z’-factor is a dimensionless parameter and is calculated with the formula below:

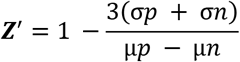

Where **3σ**_+_ is 3 times the standard deviation of the positive control, **3σ**_-_ is 3 standard deviations from the negative control, **μ**_+_ average signal of the positive control and **μ**_-_ average signal of the negative control. To calculate the Z’-Factor we used the values obtained from wells treated with pCMBs as a negative control and wells treated with classical mitochondrial toxicants and pCMBs as a positive control.

## Results

### Mitochondrial Toxicants Produced an Acute Increase of Intracellular Lactate Levels

The mitochondrial poison azide, a well know ETC inhibitor at the cytochrome c level, induced an acute increase of lactate accumulation in astrocytes, brain cells that are highly sensitive to mitochondrial toxicants (San Martin et al. 2013). This observation is explained by blockage of mitochondrial pyruvate consumption and secondary glycolytic activation. Thus, accumulated cytosolic pyruvate is converted to lactate producing its cytosolic accumulation within seconds. With the aim of test if azide induced-lactate accumulation can be reproduced using another mitochondrial toxicant with a different molecular target, we performed single-cell experiments exposing primary culture of astrocytes from the mouse cortex expressing Laconic (**Figure 1A**) to different mitochondria inhibitors that block the ETC at different levels (**Figure 1C**). To take advantage of the maximal dynamic range of the lactate FRET sensor the baseline measurements were performed using buffer without carbon sources to force start the experiments with lactate levels below 100 μM. Treatment with rotenone a complex I inhibitor, antimycin a complex III inhibitor, azide a cytochrome c inhibitor, and oligomicyn an ATPase blocker, induced an acute cytosolic lactate accumulation in astrocytes (**Figure 1B**). Intracellular lactate levels rapidly increased and reached a new steady-state in seconds, indicating fast mitochondrial dysfunction. Therefore, toxicity-induced lactate accumulation can be triggered by the blockage of ETC at different points. These observations suggest that intra-cellular lactate levels can be used as a wide specificity readout of mitochondrial dysfunction.

**Figure 1.**
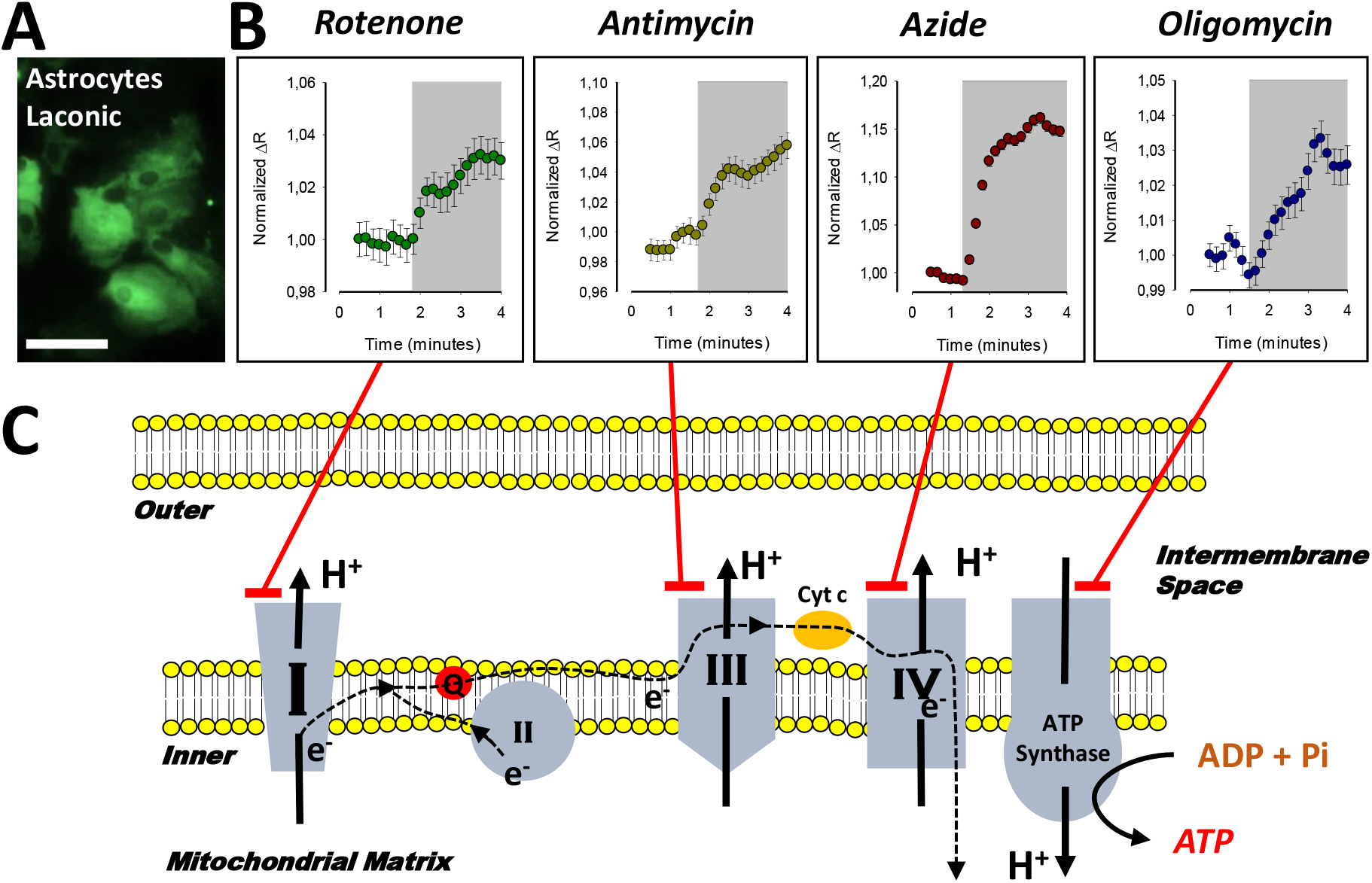
Modulation of Astrocytic Lactate Metabolism by Mitochondrial Toxicants. A) Primary culture of cortical astrocytes from mice imaged at 440 nm excitation/535 nm emission B) Cytosolic lactate accumulation induced by treatment with 32 μM rotenone, 16 μM of antimycin, 5 mM azide and 80 μM of oligomycin. Each data point is from an average of ten cells from three independent experiments. C) Schematic representation of molecular targets for each toxicant used in the primary culture of astrocytes.

### Oxidative Reporter Cell Line Generation

Sensitive cell-based system to detect mitochondrial toxicity requires a highly activated oxidative metabolism. Primary cultures, in contrast to cell lines, are highly oxidative due to their non-cancer origin, therefore they are more sensitive to mitochondrial toxicants (Wills et al. 2015). However, clonal cell lines offer advantages over primary cultures due to their HTS applications, such as low-cost production/maintenance, straightforward procedures for sample preparation, and they do not require the use of animals. in spite the fact that immortalized cell lines generate most of their ATP from glycolysis (Vander Heiden et al. 2009), there are strategies to boost their oxidative metabolism. For instance: i) forcing cells to grow in media with galactose as the only carbon-source (Marroquin et al. 2007), ii) use growth media with oleic acid as the only carbon source (Andreux et al. 2014) and iii) perform the experiments with cells cultured at a high confluence to increase cell-to-cell contact and to decrease the proliferative phenotype (Andreux et al. 2014). Therefore, we determined the magnitude of glycolytic and oxidative metabolism of a panel of cell lines and primary cultures of astrocytes using Warburg Index (WI) (**Supplementary Figure 1A**). This parameter allows glycolytic and oxidative metabolism to be evaluated at single-cell resolution (San Martin et al. 2013). The protocol consists in the acute reversible perturbation of the intracellular lactate steady-state levels blocking the ETC with azide and then promoting its accumulation to inhibit lactate exit through the MCT. During the first seconds the azide slope is a readout of pyruvate consumption and therefore mitochondrial metabolism and pCMBs promotes lactate accumulation, which is proportional to its production through glycolysis (**Supplementary Figure 1B**). Within the panel of cell lines, we tested: epithelial HEK293 cells, tumoral cell line HeLa, primary astrocytes from mouse cortex, hepatic cell line HepG2, and tumoral cell line MDA-MB-231. Cells were cultured at 100% confluency with their recommended media. MDA-MB-231 cells were grown using Leibowitz media with galactose as the only carbon source to boost oxidative metabolism.

Cell distribution based on their pCMBs/Azide slopes indicates that MDA-MB-231 are as oxidative as astrocytes, since the major part of both cell types are in the right lower quadrant (**Supplementary Figure 1C**), which is an oxidative zone characterized by higher azide slopes than pCMBs. In agreement with previous results, primary culture of astrocytes present WI below one (San Martin et al. 2013), since the WI is the glycolytic/oxidative metabolism quotient, this value confirms that astrocytes are highly oxidative. Using WI from astrocytes as a reference, we determined this parameter for tumoral and epithelia cell lines to find an oxidative system to substitute primary culture of astrocytes. HEK293 cells shown an intermediate WI value (**Supplementary Figure 1D**), which is in agreement with its epithelial origin and is consistent with previous results (San Martin et al. 2013). Within the tumoral cell lines, HeLa and HepG2 presented high and variable WI values, which is expected due their tumoral origin (**Supplementary Figure 1D**). However, the breast adenocarcinoma cell line MDA-MB-231 presents low WI, resembling astrocytes values (**Supplementary Figure 1D**), indicating that MDA-MB-231 presents a highly activated oxidative metabolism in the presence of galactose as the only carbon source and potentially this cell line could be highly sensitive to mitochondrial toxicants. Our results using WI to evaluate cellular metabolism of a panel of cell types, show that the tumoral cellular line MDA-MB-231 grown using galactose as the only carbon source, is a good alternative model for evaluating mitochondrial toxicants.

To generate a cell system with a high signal-to-noise ratio suitable for standard multiplate readers, we developed a clonal cell line with high levels and homogeneous Laconic expression. To do so, we produced a lentivirus particle to infect and transduce Laconic in oxidative MDA-MB-231 cells. From pooled cells with diverse levels of Laconic expression we selected high expression clones through FACS (<u>F</u>luorescent <u>A</u>ctivated <u>C</u>ell <u>S</u>orting). Within the high expression clones, we selected over-expressing Laconic clonal MDA-MB-231 cells that showed a homogenous cytosolic and nuclear exclusion expression signal (**Figure 2A**). We termed this cell line “MitoTox Reporter”. Exposure of these cells to classical mitochondrial poisons produced a robust increase in the fluorescent signal, indicating acute toxicity-induced lactate accumulation. For instance, treatment with rotenone, antimycin, oligomycin, and myxothiazol produced an acute, fast, and robust increase of intracellular lactate levels (**Figure 2B, C, D and E**). The response was higher in comparison with the increase of lactate levels induced by mitochondrial toxicants in astrocytes, demonstrating high sensitivity to mitochondrial toxicants. With this evidence in hand, cells were tested in a multi-well plate reader.

**Figure 2.**
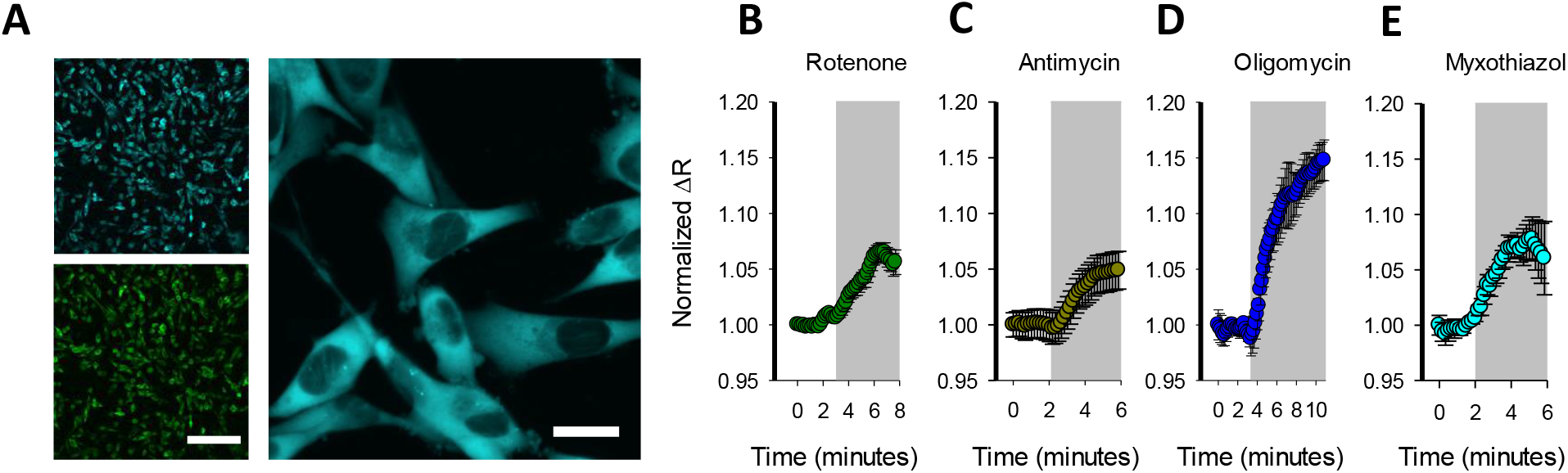
Characterization of “MitoTox Reporter” Cell Line. A) Confocal images from FACS selected clonal MDA-MB-231 cells expressing Laconic. Size bar for 20x and 60x represent 100 μm and 10 μm, respectively. Cytosolic lactate accumulation induced by treatment with: B) 32 μM rotenone, C) 16 μM of antimycin, D) 80 μM of oligomycin, and 10 μM myxothiazol. Each trace is the average of ten cells from one representative.

### Mitochondrial Toxicants Produced Toxicity-Induced Lactate Accumulation Detectable by a Multi-Plate Reader

Fluorescence-based readouts are amenable to HTS. To assess the detection of toxicity-induced intracellular lactate accumulation by mitochondrial toxicants in a multiwell plate format, MitoTox reporter cells were seeded in 96-well plates and incubated without CO_2_ equilibration at 37 °C until they reached a maximal confluency, using a galactose rich media Leibowitz. Cells were exposed to a high concentration of classical mitochondrial toxicants: 32 μM rotenone, 16 μM antimycin, 5 mM azide, 80 μM oligomycin, and 10 μM of myxothiazol. The effects were measured at 5, 10, 30, and 60 minutes. All these mitochondrial toxicants produced an acute lactate accumulation that was detectable in just 5 minutes at 37 °C. The amplitude of the response was variable and based on ΔR% not all the mitochondrial toxicants produced enough lactate accumulation to saturate the sensor, not making it possible to take full advantage of the sensor’s dynamic range (**Supplementary Figure 2A, B, C, D, E and F**). Also, the response was not stable over-time especially with azide which declined after 5 minutes (**Supplementary Figure 2D**). Positive control 10 mM lactate also does not induce enough lactate accumulation to saturate the sensor (**Supplementary Figure 2A**). However, some wells with higher responses were consistent with the maximal FRET change of Laconic, which is 40 %. Each experimental series included a well with the solvents used to prepare the mitochondrial toxicants in order to test if they produced interference. DMSO, ethanol, and KRH buffer did not produce any interference at the concentration used to perform the toxicity experiments, since they did not induce a significant effect on ΔR% over-time (**Supplementary Figure 2A, B, C, D, E and F**). Data dispersion was high and ΔR% were not high enough to produce a method suitable for HTS application but gives a proof of the concept that is possible to detect increments of lactate levels induced by a mitochondrial toxicant in a single-well format using a standard multiplate reader. However, the response needs to be improved to make the method suitable for HTS.

### Optimization of Toxicity-Induced Lactate Accumulation Blocking the Monocarboxylate Transporters (MCT)

To develop a suitable method for HTS application, it was necessary to increase de amplitude of toxicity-induced lactate accumulation and decrease data variation. In order to tackle these challenges, we boosted the response induced by mitochondrial dysfunction, blocking lactate exit and forcing sensor saturation by pharmacological means. To block lactate efflux, we used pCMBs (4-(Chloromercuri) benzenesulfonic acid) an inhibitor of MCTs (Wilson et al. 2005), which is a non-permeable sulfhydryl group modifier agent with broad specificity including MCT4, which is the main monocarboxylate transporter present in MDA-MB-231 cells (Baenke et al. 2015). We added 50 μM pCMBs to block lactate exit at the same time as mitochondrial toxicants such as rotenone, antimycin, azide, oligomycin, and myxothiazol. The effects of each toxicant were measured at 5, 10, 30, and 60 minutes (**Figure 3A, B, C, D, E and F**). These mitochondrial toxicants produced acute intracellular lactate accumulation in MDA-MB-231 cells tested in a 96 well plate format. The amplitude of the signal with pCMBs was higher and the data present low intra assay variability compared to the experiments without pCMBs. MCT blockage did not produce lactate accumulation, discarding possible interference with basal lactate production. This protocol provided a more robust assay with low variability and higher changes of ΔR% induced for mitochondrial toxicants in a multiplate reader, which are critical parameters to evaluate the suitability of the methodology for HTS applications. The assay was stable at least for 60 minutes for rotenone, antimycin, and myxothiazol (**Figure 3B, C, and F**), 30 minutes for azide (**Figure 3D**) and 10 minutes for oligomycin (**Figure 3E**).

**Figure 3.**
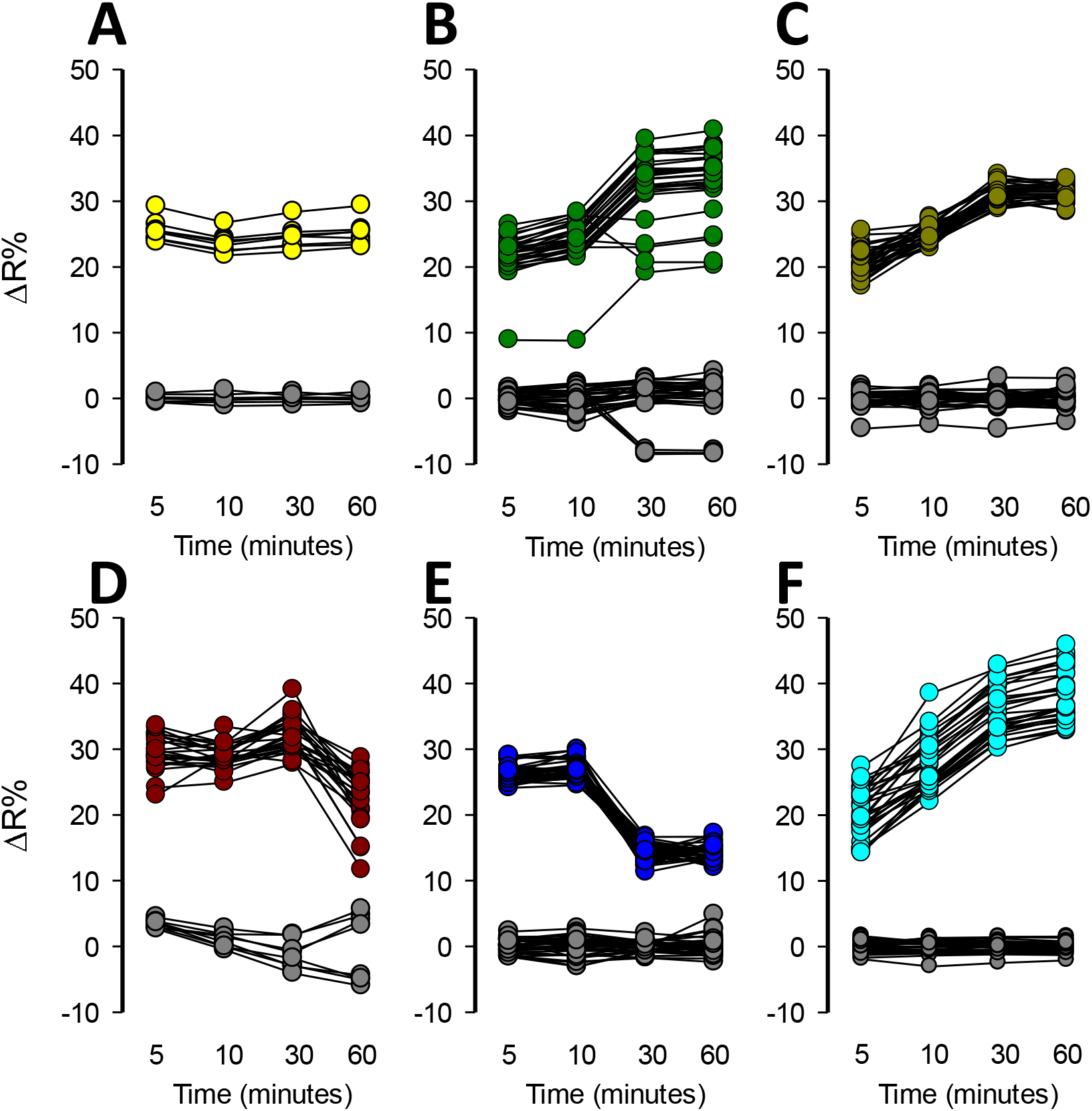
Enhancement of Lactate Accumulation by Transport-Stop Protocol. All the experiments were performed adding 50 μM of pCMBs together with the mitochondrial toxicant. Measurements were performed at 5, 10, 30, and 60 minutes. A) pCMBs control, B) 32 μM rotenone, C) 16 μM antimycin, D) 5 mM azide, E) 80 μM oligomycin and F) 10 μM myxothiazol. Solvent control (gray circles): KRH buffer for azide, 0.8% DMSO for rotenone, oligomycin, and myxothiazol and 0.16% ethanol for antimycin. All the experiments were performed at 37 °C.

To evaluate the suitability of the improved assay for HTS applications, a coefficient called the Z’-Factor was calculated (Zhang et al. 1999). This coefficient reflects the assay’s signal dynamic range and the data variation associated with the signal measurements. Z’-factor values between 0.5 – 1 mean that a hit can be identified with a confidence of 99.73 – 100% using a single-well. Values below 0.5 indicate the necessity to perform duplicates or triplicates for each molecule and values below 0 indicated the unsuitability of the assay for HTS application. We performed a series of experiments to calculate the Z’ -factor using different mitochondrial poisons and replicate plates on different experimental days. Treatment with each mitochondrial toxicant produced acute lactate accumulation with low intra-well variability and a high ΔR% increase almost reaching the reporter maximal fluorescent change. All our assays using different mitochondrial toxicants reached Z’-factor above 0.3. Majority of our replicates reached a Z’-Factor over 0.5 supporting the suitability of the assay to identify a mitochondrial toxicant in a single-well screening (**Figure 4A – L**). For all mitochondrial toxicants tested the better Z’-Factor was obtained 10 minutes after treatment. The intra and inter-plate variability for the improved method was determined calculating the coefficient of variation CV. All the mitochondrial toxicants and positive control had a low intra plate CV% below 20 and comparison between CV% values between different plates indicate low inter-plate variability (**Supplementary Figure 3**). Our date shows that the improved method using a pharmacologic approach to block the lactate exit is suitable for HTS applications.

**Figure 4.**
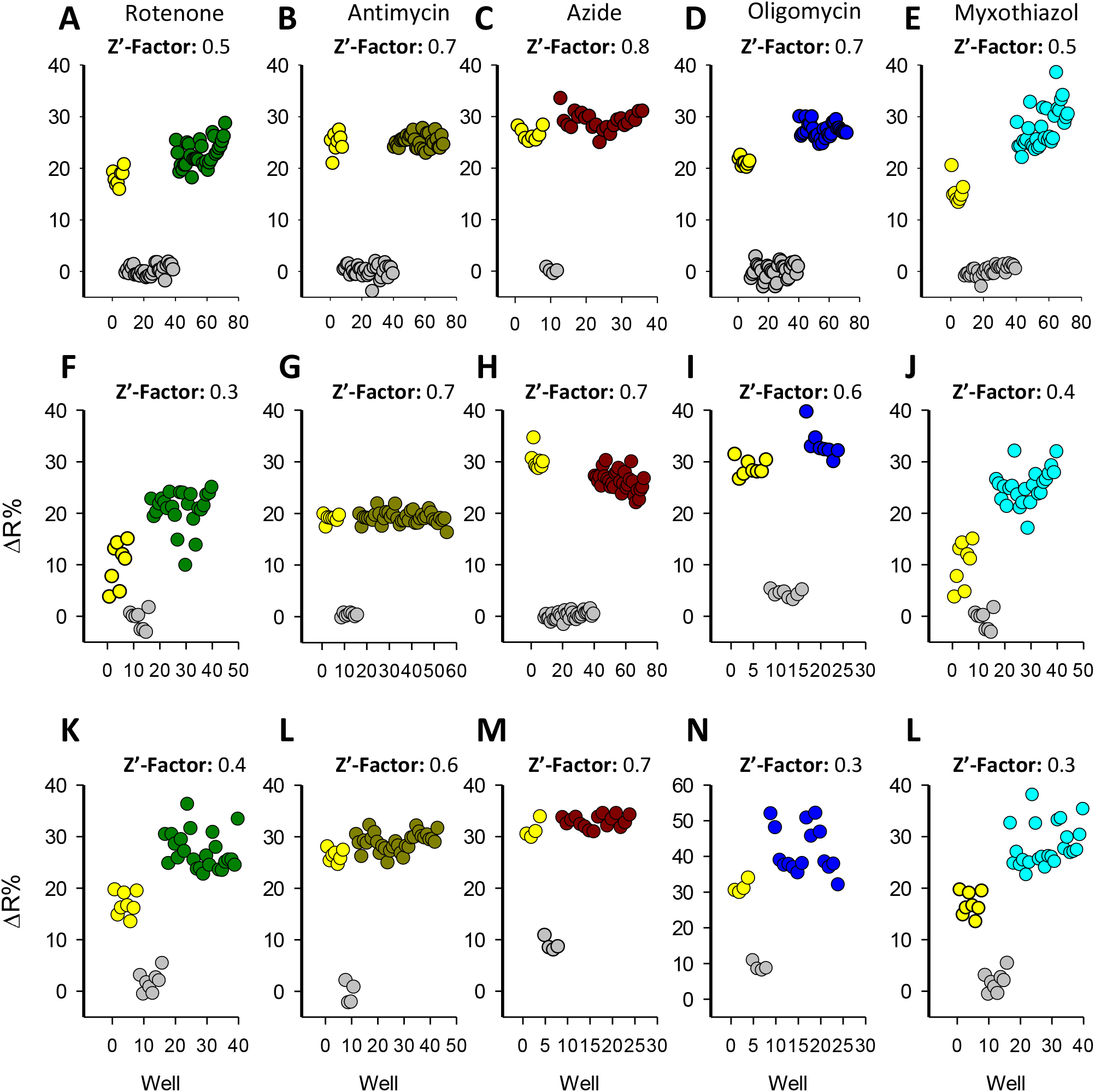
High throughput Suitability Characterization. The Z’-factor was calculated using wells treated with 32 μM rotenone, 16 μM antimycin, 5 mM azide, 80 μM oligomycin, and 10 μM myxothiazol as positive controls. Negative control wells were treated with 50 μM of pCMBs (gray circles) and nanosensor response control with 10 mM lactate (yellow circles). All the Z’-Factors were calculated at 10 minutes post-treatment. Experiments from three independent assays for each mitochondrial toxicant, performed on different experimental days.

### Sensitivity of Mitochondrial Toxicity Detection

To assess the sensitivity of our methodology we determined the IC_50_ for a panel of classical mitochondrial toxicants. Dose-response curves were constructed by treating the cells with an increasing concentration of rotenone, antimycin, azide, oligomycin, and myxothiazol. We measured the toxicity-induced lactate accumulation at 30 minutes of treatment. To quantitatively evaluate the potency of these compounds, we fitted our data to a rectangular hyperbola to calculate the IC_50_ for each toxicant. We determine that the complex III inhibitor antimycin was the most potent mitochondrial toxicant from our panel with an IC_50_ of 0.24 ± 0.48 nM (**Figure 4B**), followed by complex I inhibitor rotenone which has an IC_50_ 2.7 ± 1.41 nM (**Figure 4A**). Complex III myxothiazol reached an IC_50_ of 1.2 ± 0.55 μM (**Figure 4E**) and the ATPase inhibitor oligomycin reached IC_50_ of 38 ± 15 nM (**Figure 4D**). The weakest molecule of our mitochondrial toxicants panel was azide, a cytochrome c inhibitor which reached an IC_50_ of 1.44 ± 0.76 mM (**Figure 4B**). To gain a wider view about the sensitivity of our method, we confronted the IC_50_ from our methodology with the state-of-the-art methods to assess mitochondrial dysfunction (**Supplementary Table 1**). Our methodology allows the detection of mitochondrial toxicity induced by classical toxicants faster than ATP, MMP, GSH, viability and ROS measurements and presents improved sensitivity compared to OCR and ECAR when detecting mitochondrial dysfunction induced by rotenone, antimycin, and oligomycin using Seahorse^®^ technology. These results suggest that real-time monitoring of intracellular lactate levels is more sensitive and faster than the state-of-the-art technology to evaluate mitochondrial dysfunction using standard multiplate readers.

### Pilot Screening

The capability of the assay to identify previously described toxic drugs was explored through a pilot screening assay. We selected 13 compounds from a variety of therapeutic classes to perform a pilot screening assay using MitoTox Reporter cells and a standard multiplate reader. Among the selected compounds we included thiazolidinediones such as ciglitazone, troglitazone, and rosiglitazone which are antidiabetic agents with known organ toxicity (Hernandez et al. 2011), camptothecin a topoisomerase inhibitor that produces hepatic toxicity (O’Leary and Muggia 1998), prazosin an alpha-adrenergic antagonist with no reported associated toxicity, simvastatin a long-established hydroxy-methylglutaryl coenzyme A (HMG-CoA) reductase inhibitor safe if administered alone (Pedersen and Tobert 2004), metformin a safe antidiabetic agent (van der Aa et al. 2016), terfenadine a worldwide withdrawn antihistamine due to cardiac arrhythmia (Smith 1994), a nonsteroidal antiandrogens nilutamide, and flutamide which are used for the treatment of prostate cancer and produced a decrease of oxygen consumption and hepatotoxicity (Berson et al. 1994; Giorgetti et al. 2017), etoposide semisynthetic derivative of podophyllotoxin with broadly anticancer activity with low side effects (Kobayashi and Ratain 1994), acetylsalicylic acid and lidocaine both with no reported cell toxicity effects. To perform the assays, we prepared master stocks for each compound in 0.2% DMSO. Assays were performed with compounds at final concentrations of 1, 5, and 10 μM and the response of MitoTox Reporter cells were measured at 5, 10, 30, and 60 minutes. Thiazolidinediones such as ciglitazone, troglitazone, and rosiglitazone produces a robust increase over 14% of ΔR at 10 μM, which is consistent with an acute increase of cytosolic lactate due to mitochondrial dysfunction. The effect was also detected at 5 μM for troglitazone (**Figure 5A and B**). Lactate accumulation was stable during 60 minutes after drug treatment at 10 μM of each compound (**Supplementary Figure 4**). These results are consistent with the known cellular toxicity identified for these class of molecules (Nanjan et al. 2018). Additionally, worldwide withdrawn antihistamine terfenadine produced a rapid and robust increase of 15.64 ± 1.32 of ΔR% at 10 μM, without any measurable effects at 1 or 5 μM (**Figure 5A and B**). Lactate accumulation was stable during 60 minutes after drug treatment at 10 μM (**Supplementary Figure 4**). Strikingly, a highly toxic anti-cancer drug camptothecin produced a potent increase of 49.18 ± 7.4 ΔR% at 10 μM consistent with high lactate accumulation into the cytosol (**Figure 5A and B**). Taking into account the reported dynamic range and sensitivity of Laconic (San Martin et al. 2013), camptothecin produced lactate accumulation over 10 mM since the drug treatment induced a maximal change in the dynamic range of the sensor. Also, a potent effect at 5 μM was observed, but no change in ΔR% was detected at 1 μM of the drug. The effects on lactate metabolism were stable during 60 minutes after drug treatment at 10 μM (**Supplementary Figure 4**). Also, nilutamide and flutamine with known hepatotoxic effects, showed an opposite effect on lactate metabolism. Although nilutamine did not produce any effects flutamine induced an 8.71 ± 2.06 ΔR% at 10 μM. As a negative-control we selected a compound with no reported side-effects. Among the selected molecules we had metformin, prazosin, simvastatin, etoposide, aspirin, and lidocaine. None of these molecules produced a lactate accumulation at 1, 5, and 10 μM (**Figure 5A and B**). These results validate our methods to identify molecules with a toxic profile.

**Figure 5.**
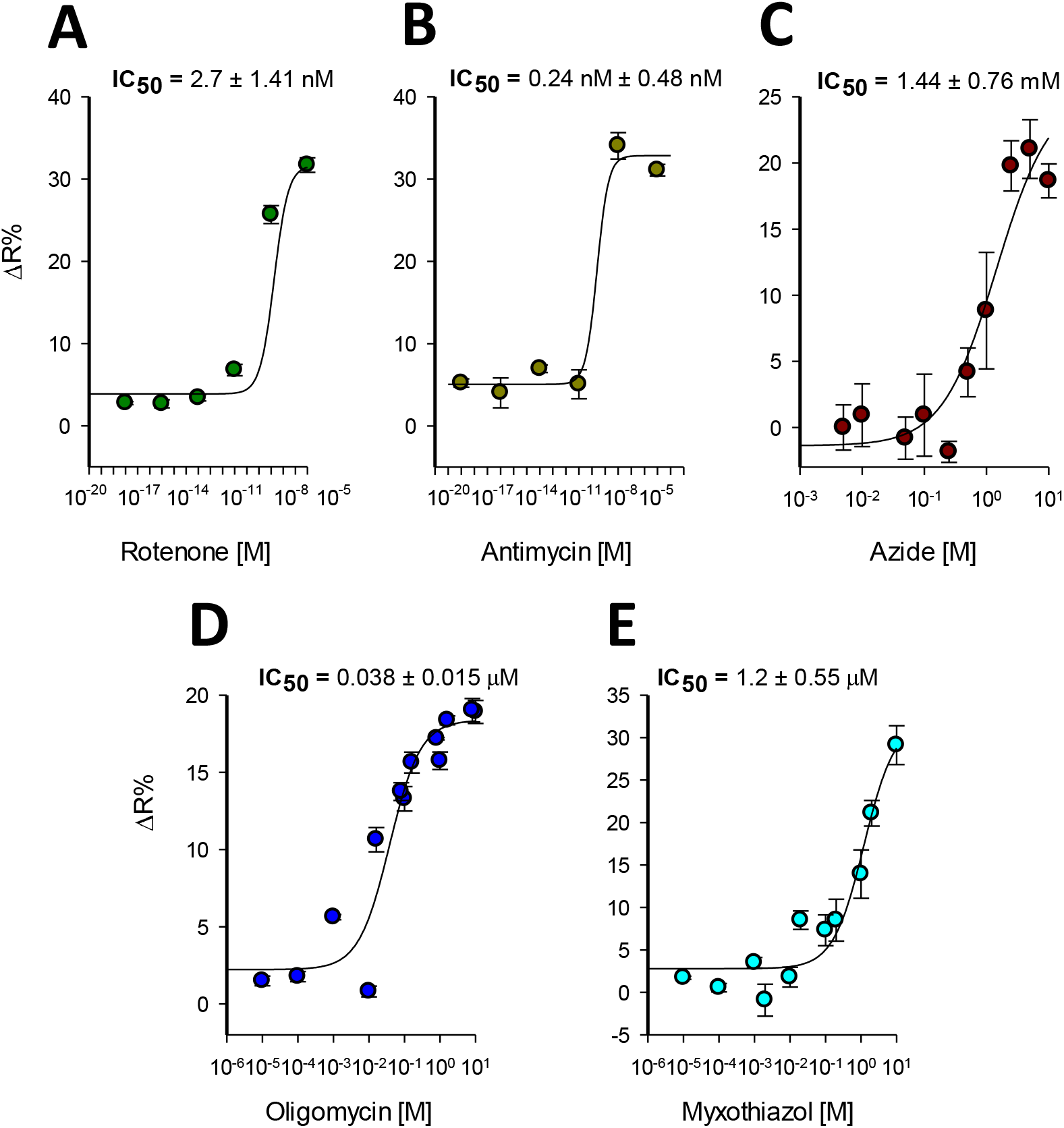
IC_50_ for Classical Mitochondrial Toxicants. Dose-response Lactate accumulation induced by classical mitochondrial toxicants was measured at: A) 10×10^-5^, 10×10^-8^, 10×10^-11^, 10×10^-14^, 10×10^-17^ and 10×10^-20^ in molar of rotenone and B) antimycin C) 10, 5, 2.5, 1, 0.5, 0.25, 0.1, 0.05, 0.01 and 0.005 in mM of azide D) 16, 10, 8, oligomycin and E) 10, 2, 1, 0.2, 0.1, 0.02, 0.01, 0.002, 0.001, 0.0001 and 0.00001 in μM of myxothiazol. Data was collected at 10 minutes after toxicant treatment at 37°C. Data is the average from three independent experiments.

**Figure 6.**
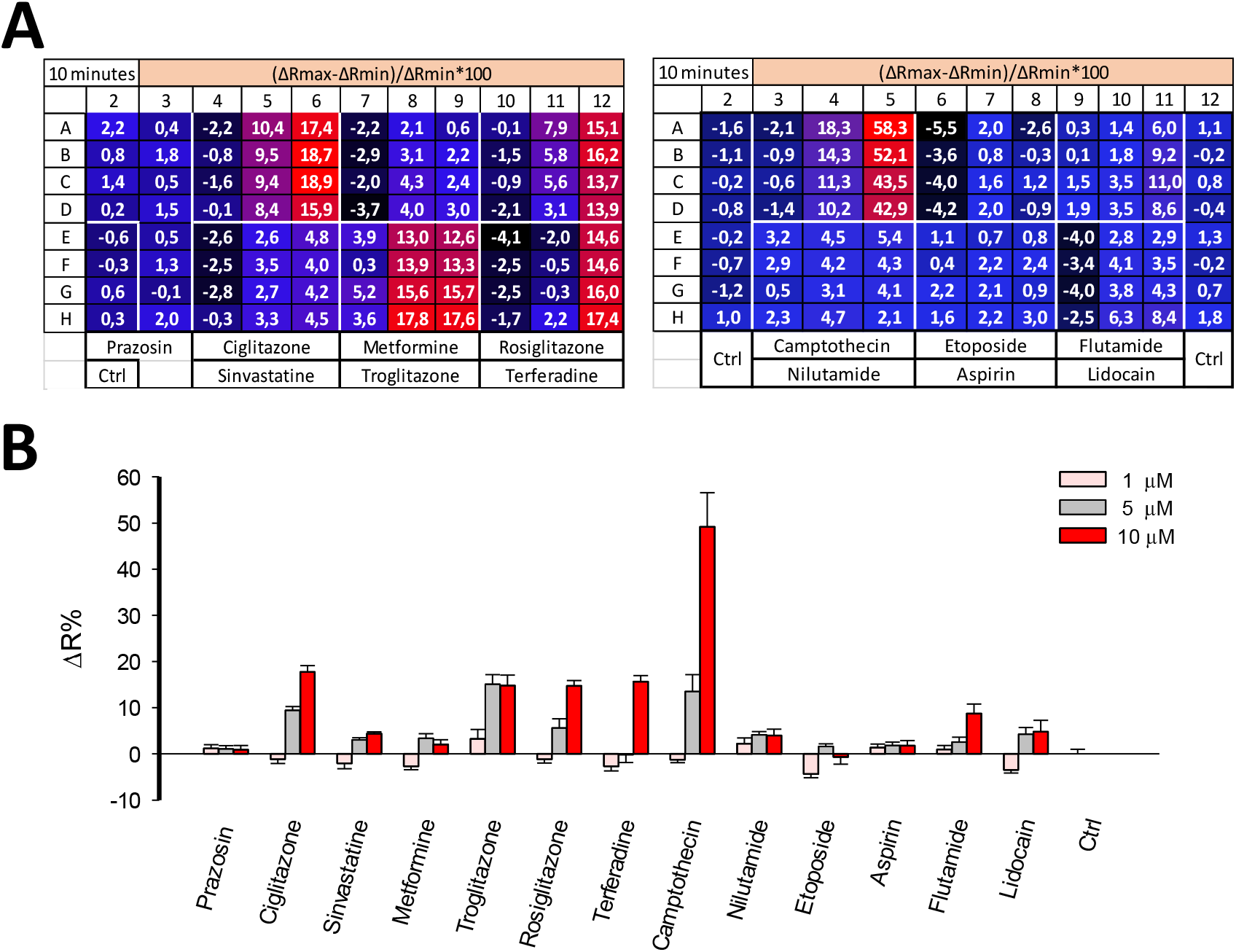
Pilot Screening. A) Schematic representation of the 96 well plate format with their corresponding ΔR% values in pseudo color codification. Red and blue colors represent high and low ΔR%, respectively. Drugs were used at 1, 5 and 10 μM and effects were measured after 30 minutes of treatment at 37 °C B) Bar plot of the effect on lactate levels of a panel of drugs at 1, 5, and 10 μM at 37°C after of 30 minutes of pharmacological treatment. Data represent the average of 4 replicate wells of one representative experiment.

## Discussion

This paper describes a phenotypic high-throughput screening method for the identification of mitochondrial toxicants from chemical libraries. The methodology takes advantage of toxicity-induced intracellular lactate accumulation monitored using a genetically-encoded lactate indicator, Laconic. To test the methodology, lactate accumulation was induced by acute pharmacological blockage of ETC at different levels by classical mitochondrial toxicants. The suitability for HTS applications was tested and sensitivity was evaluated with the current state-of-the-art approaches to assess mitochondrial dysfunction. Pilot screening was performed to evaluate the capabilities of the methodology to identify previously described toxic drugs.

Our results from IC_50_ determination using classical mitochondrial toxicants show that is more sensitive and requires less time for readout measurements. Indeed, our method detect mitochondrial dysfunction using nanomolar concentrations of the toxicant rotenone in just 30 minutes, much better than current techniques that require hours of pre-treatments. This improved sensitivity suggests that our methodology could be useful for detecting subtle changes in mitochondrial physiology, which are responsible for the appearance of side-effects well after drug approval during the commercialization phase. Additionally, our pilot screening showed the capability of the methodology to detected previously described mitochondrial toxicants with improved sensitivity. For instance, the antidiabetic agents thiazolidinediones induced a robust 20% ΔR change at 10 μM that covers half of the sensor’s dynamic range. Previous studies using Seahorse^®^ technology show that concentrations over 100 μM of ciglitazone and troglitazone are required to change the OCR and ECAR of HepG2 cells by 50% and no effects were detected between 0.13 and 100 μM for rosiglitazone (Nadanaciva et al. 2012). Measurements of MMP showed that the IC_50_ for ciglitazone, rosiglitazone, and troglitazone are 135.7, >1000 and 175.6 μM, respectively (Li et al. 2014). The antihistamine drug terfenadine induced a 20% ΔR change at 10 μM, meanwhile MMP measurements had an IC_50_ of 4.4 (Li et al. 2014). The highly toxic anti-cancer drug camptothecin produced a potent increase of 49.18 ± 7.4 ΔR% at 10 μM and 13.52 ± 3.6 at 5 μM. Spite of their known cellular toxicity, this robust response in intra-cellular lactate accumulation over 10 mM is not obvious, since MMP measurements determine the IC_50_ of 205.6 μM (Li et al. 2014). Our negative control molecules metformin, etoposide, aspirin, and lidocaine, which are molecules without reported cellular toxicity, did not induce intra-cellular lactate accumulation, in agreement with previous results (Li et al. 2014). The exceptions were prazosin and simvastatin, which were not detected by our methods, but they induced MMP depolarization with an IC_50_ of 6.3 and 63.2 μM, respectively (Li et al. 2014). These results can be considered false positive hits from MMP measurements, because they are considered to be relatively safe drugs (Molgaard et al. 1991). Also, nilutamide did not produce ΔR% changes at all the concentrations tested, despite of its known toxicity profile (Berson et al. 1994), being a false-negative result. A possible explanation for this behavior is that the drug stops pyruvate consumption, but without secondary glycolytic activation, so the amount of lactate is not enough to produce saturation of the sensor. In this case we expect to improve the sensitivity using a version of Laconic with higher affinity in the micromolar range, thus lactate increase will induce a fast sensor saturation. These results indicate that our methodology is complementary with current methods to evaluate mitochondrial function such as MMP measurements, therefore we consider that potential hit should be validated with complementary techniques.

The introduced methodology based on a FRET lactate sensor presents limitations. We obtained Z’-Factors > 0.5, since the maximal amplitude of fluorescent change that Laconic can afford is 40% and the intra-plate variability is < 20%, we are within the limit of an appropriate Z’ Factor for HTS applications. Therefore, if the data variability undergoes a slight increase this will produce a major impact on the Z’ Factor, making it difficult to obtain values > 0.5 and the method will not suitable for HTS applications. Another limitation is the steady-state fluorescent measurements are sensitive to color interferences, therefore colored compounds cannot be evaluated using this methodology and they must be discarded previously to avoid false positive/negative hits.

Phenotypic and convergent readouts are required to evaluate cellular toxicity at the beginning of the drug development process to decrease the attrition rate in the pharmaceutical industry. Genetically-encoded indicators are emerging tools that allow cell physiology in intact systems to be evaluated. We envisage that the continuous development and optimization of sensors for metabolites, focusing on improvements in the dynamic range, brightness, and affinity for HTS purposes, will allow the development of a robust phenotypic cell-based assay for the pharmaceutical industry.

## Acknowledgements

This work was partly supported by the CORFO project “High Technology Business Innovation” 14IEAT-28662 and CONICYT through FONDECYT Initiation into Research 11150930 program. The Centro de Estudios Científicos (CECs) is funded by the Chilean Government through the Centers of Excellence Basal Financing Program of CONICYT.

## Author Contribution Statement

Y.C.B performed the experiments in 96 well plates. S.C and R.A.M performed the experiments using primary astrocytes. P.Y.S generated lentivirus particles and MitoTox reporter cells. K.A cell culture preparation. L.F.B and A.S.M conceived the project and wrote the manuscript.

