## Supplementary Information for "MitoToxy Assay: a novel cell-based method for the assessment of metabolic toxicity in a multiwell plate format using a lactate FRET nanosensor, Laconic"

^1^Centro de Estudios Científicos (CECs), Postal Code 5110466, Valdivia, Chile.

^2^Universidad Austral de Chile (UACh), Postal Code 5110566, Valdivia, Chile.

**Corresponding author:**

Alejandro San Martín, Centro de Estudios Científicos (CECs), Avenida Arturo Prat 514, Postal Code 5110466, Valdivia, Chile.

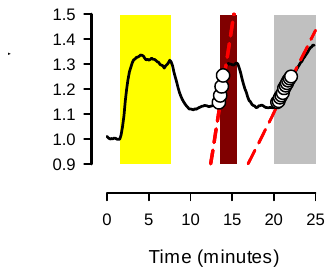

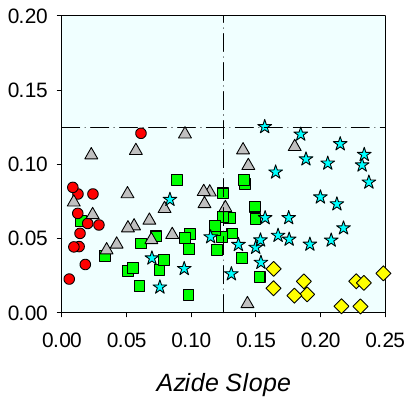

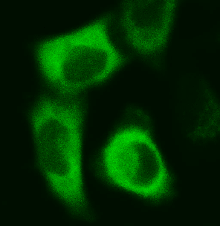

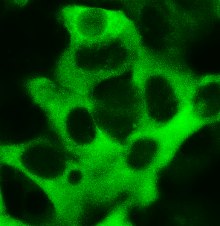

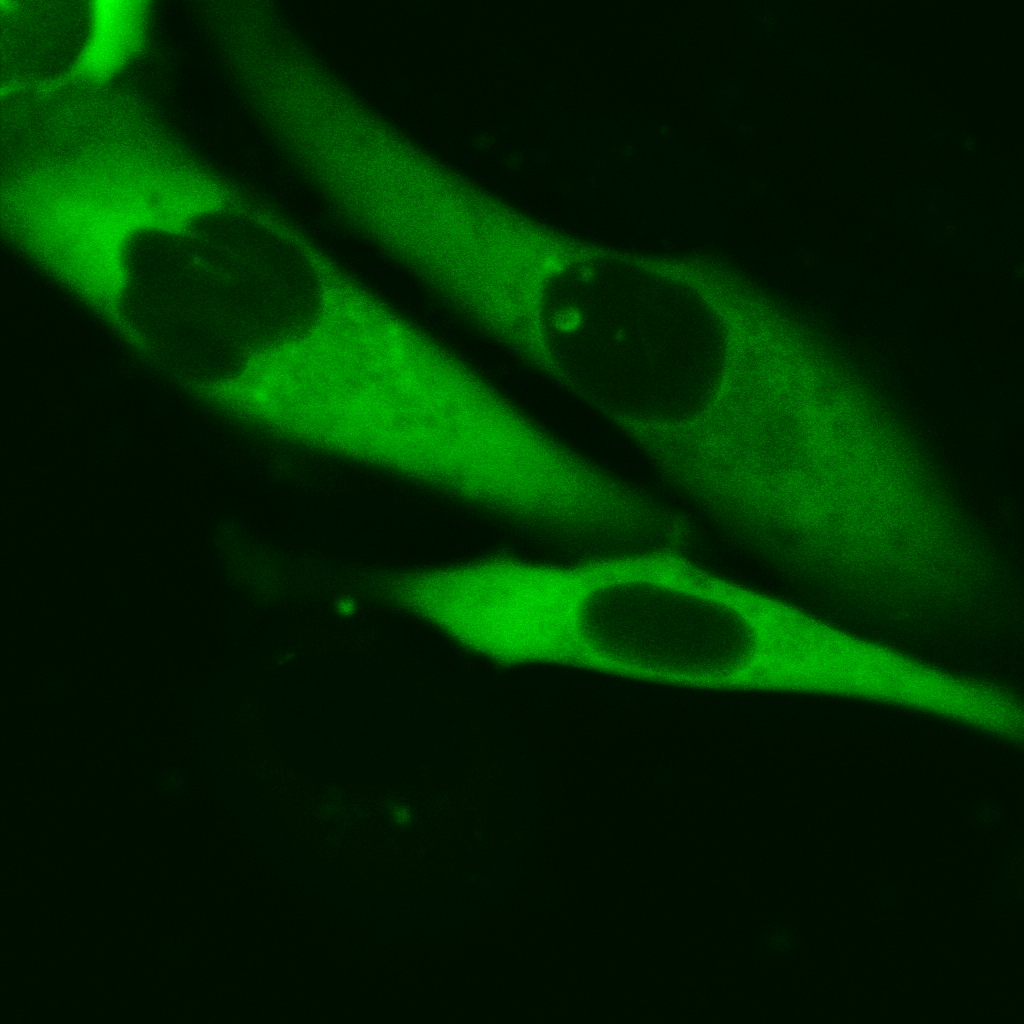

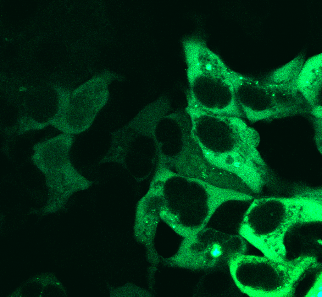

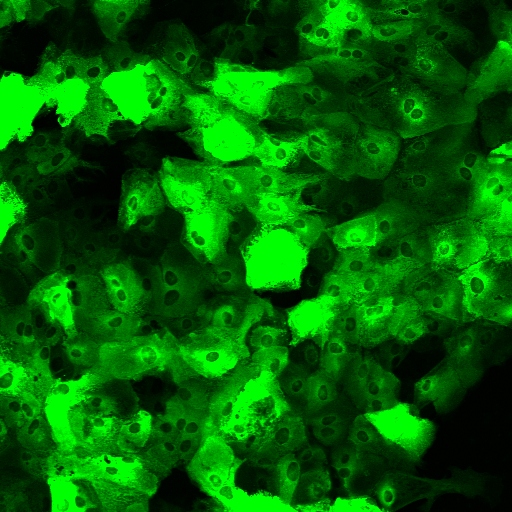

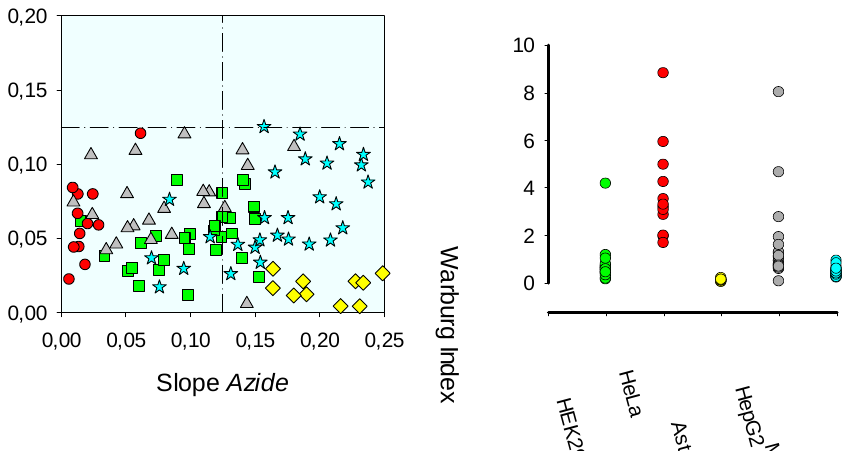

**Supplementary Figure 1. Selection of an Oxidative Cell Line.** A) A panel of epithelial, tumoral cellular lines, and primary astrocytes expressing cytosolic Laconic. Images were taken by confocal microscopy. Scale bar 10 µm for HEK293, HeLa, HepG2 and MDA-MB-231. Scale bar 50 µm for astrocytes. B) Standard protocol for WI determination. Oxidative and glycolytic metabolism were explored in a primary culture of astrocytes by ETC inhibition with 5 mM Azide and stopping lactate transport with 250 µM, respectively. C) Response distribution of pCMBs slope (glycolysis) and azide (OXPHOS). Oxidative cells are in the right lower quadrant. D) WI calculations for each cell using the quotient from pCMBs/azide slopes. Data from three independent experiments.

**HeLa**

**HepG2**

**MDA-MB-231**

**Astrocytes**

**HEK293**

**A**

**C**

**D**

**B**

Azide

pCMBs

Lactate

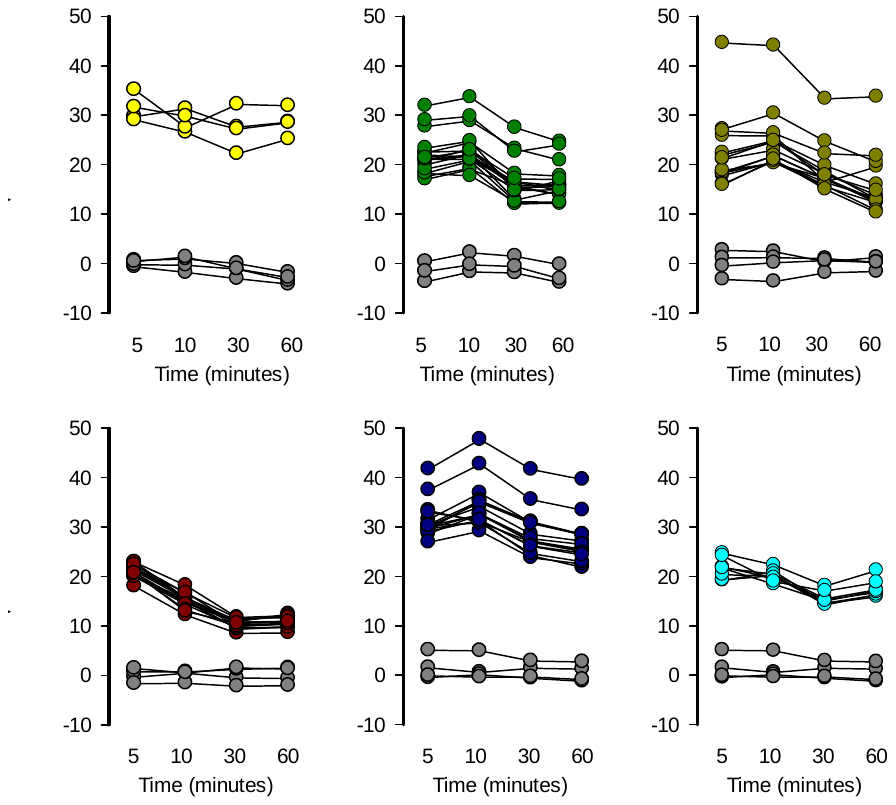

**F**

**E**

**D**

**C**

**B**

**A**

**Supplementary Figure 2. Single-well Detection of Lactate Accumulation Induced by Mitochondrial Toxicants.** Mitochondrial dysfunction induced by mitochondrial toxicants was detected in MitoTox Reporter cells in 96 well plates. Measurements using a standard multiplate reader were performed at 5, 10, 30, and 60 minutes. A) 10 mM Lactate, B) 32 µM rotenone, C) 16 µM antimycin, D) 5 mM azide, E) 80 µM oligomycin and F) 10 µM myxothiazol. Solvent control (gray circles): KRH buffer for azide, 0.8% DMSO for rotenone, oligomycin, and myxothiazol and 0.16% ethanol for antimycin. All the experiments were performed at 37 ºC.

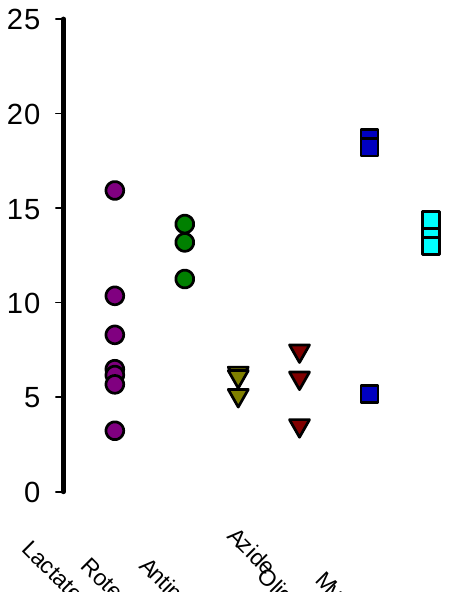

**Supplementary Figure 3. Inter and Intra-well Variability Analysis.** CV% values were calculated using the average wells from three independent plates.

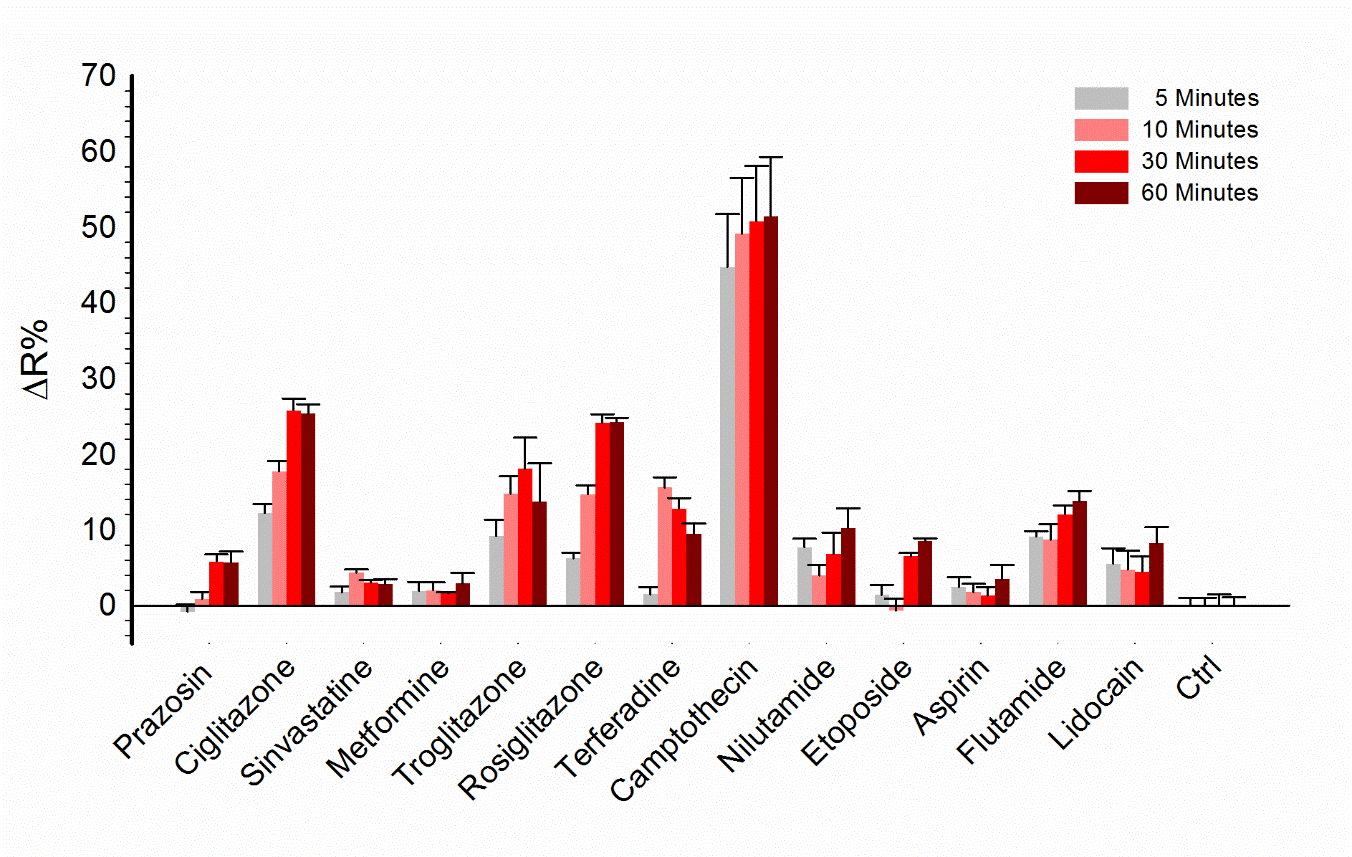

**Supplementary Figure 4. Stability of Lactate accumulation.** Lactate accumulation at 5, 10, 30, and 60-minutes of treatment with a panel of drugs at 37 ºC. Data obtained from treatment with 10 µM of each compound.

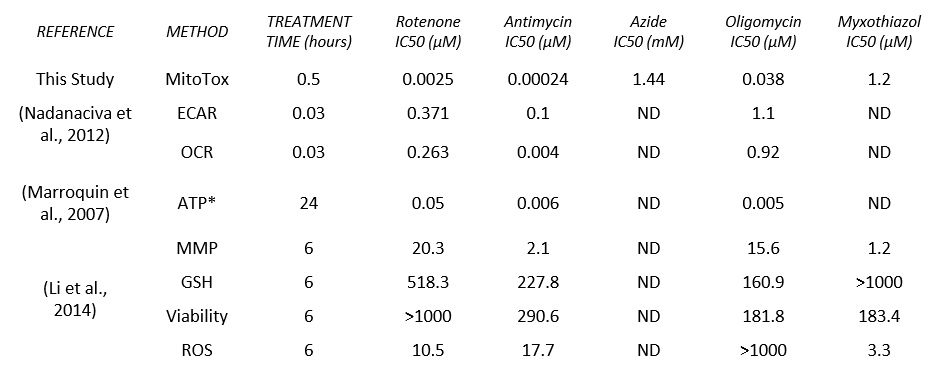

**Supplementary Table 1. IC_50_ Comparison of State-of-Art Methods to Evaluate Mitochondrial Dysfunction.** Benchmarking analysis comparing the IC_50_ archived with the Laconic based method and current technology to assess mitochondrial dysfunction.

*Approximated IC50 directly from plotted data in the original paper

ND: No Determined
